# The role of alpha power in the suppression of anticipated distractors during verbal working memory

**DOI:** 10.1101/2020.07.16.207738

**Authors:** Sabrina Sghirripa, Lynton Graetz, Ashley Merkin, Nigel C Rogasch, Michael C Ridding, John G Semmler, Mitchell R Goldsworthy

## Abstract

As working memory (WM) is limited in capacity, it is important to direct neural resources towards processing task-relevant information while ignoring distractors. Neural oscillations in the alpha frequency band (8-12 Hz) have been suggested to play a role in the inhibition of task-irrelevant information during WM, although results are mixed, possibly due to differences in the type of WM task employed. Here, we examined the role of alpha power in inhibition of anticipated distractors of varying strength using a modified Sternberg task where the encoding and retention periods were temporally separated. We recorded EEG while 20 young adults completed the task and found: 1) slower reaction times in strong distractor trials compared to weak distractor trials; 2) increased alpha power in posterior regions from baseline prior to presentation of a distractor regardless of condition; and 3) no differences in alpha power between strong and weak distractor conditions. Our results suggest that parieto-occipital alpha power is increased prior to a distractor. However we could not find evidence that alpha power is further modulated by distractor strength.

## Introduction

Verbal working memory (WM) refers to the ability to temporarily maintain and/or manipulate verbal information to guide immediate cognitive processing (Baddeley, 1992). WM consists of three stages: encoding, which involves the ‘loading’ of information into WM, which is then stored and refreshed throughout a retention period, before the information is retrieved to perform a goal-directed action (Baddeley, 1992). WM, however, is limited in capacity, highlighting the need for efficient use of WM storage by encoding task-relevant information while ignoring irrelevant or distracting information. Though the successful filtering of distractors is required for successful WM performance (Vogel and Machizawa, 2004), the neural mechanisms underlying this process are not fully understood.

Neural oscillations in the alpha frequency range (8-12Hz) are thought to play a role in WM, though the direction and magnitude of alpha modulation is task dependent. In verbal WM tasks where the encoding stimuli are presented simultaneously, alpha power during the retention period tends to increase with WM load (Jensen et al., 2002; Proskovec et al., 2019). This increase in alpha power is thought to represent a sensory gating mechanism, reflecting inhibition of the visual cortex to prevent disruption to WM maintenance occurring in frontal and parietal brain areas (Jensen and Mazaheri, 2010). However, in N-back style tasks where encoding stimuli are presented sequentially, alpha suppression during the retention period occurs with increasing load (Gevins et al., 1997; Pesonen et al., 2007; Stipacek et al., 2003). Therefore, the variation in alpha modulation during verbal WM tasks may be due to the method in which encoding stimuli are presented. To investigate this, Okuhata et al. (2013) compared the presentation mode of encoding stimuli in a Sternberg task (either simultaneous or sequential), and found that alpha power during the retention period decreased in the simultaneous task, but increased during the sequential task when compared to baseline, despite each condition being matched for WM load, providing evidence against the role of alpha activity as a sensory gating mechanism during WM.

Due to the role of alpha activity in sensory gating, alpha has also been implicated in distractor inhibition during WM. Evidence linking alpha oscillations to distractor inhibition was derived from lateralised visual WM tasks, where subjects attend to and memorise the information in a cued hemifield and ignore the information in the un-cued hemifield (i.e. the un-cued hemifield acts as a distractor during the encoding stage). In these paradigms, visual alpha activity tends to increase in the task-irrelevant hemisphere, suggesting a role of alpha power in the suppression of distracting or task-irrelevant information (Sauseng et al., 2009). Similar to verbal WM literature, results in this field are conflicting and potentially task dependent. For example, another study manipulating distractor strength in a lateralised visual WM task found that alpha power in parieto-occipital brain regions decreased in the presence of strong distractors when the distractor was present during the entire retention interval (Schroeder et al., 2018).

Less attention has been directed to investigating the role of alpha oscillations in distractor inhibition during verbal WM. Using a Sternberg task with sequentially presented memory sets and weak or strong distractors during the retention interval, Bonnefond and Jensen (2012) showed that anticipation of a strong distractor was associated with greater alpha power prior to distractor onset, and this effect correlated with increased task-performance in the presence of strong distractors. However, given that alpha oscillatory activity during WM is task dependent, it is currently unknown whether anticipating a distractor during the WM retention period leads to a modulation of alpha power in a task where the memory set is presented simultaneously.

To address whether alpha power is modulated by distractor strength, we employed a modified Sternberg task where the memory set was presented simultaneously, and then displayed a strong or weak distractor during the retention period. We analysed spectral power in the lead up to the distractor in the retention period to examine whether alpha activity reflected anticipation of the strength of a distractor. Our main hypothesis was that alpha power would show a larger increase before the onset of a strong, relative to a weak distractor.

## Method

### Participants

20 healthy, right-handed, young adults participated in the study (mean age = 22.8 years, SD = 4.09 years, 11 female). Exclusion criteria involved a history of neurological/psychiatric disease, use of central nervous system altering medications, history of alcohol/substance abuse and uncorrected hearing/visual impairment. All participants gave informed written consent before the commencement of the study, and the experiment was approved by the University of Adelaide Human Research Ethics Committee.

### Modified Sternberg Task

The modified Sternberg WM task used stimuli presented by PsychoPy software (Peirce, 2007) (figure 1A). At the beginning of each trial, the participant fixated on a cross in the centre of the screen for 2 s. A memory set consisting of 5 consonants was then shown for 1 s, followed by a 4 s retention period. 2 s into the maintenance period, a weak (3 hash symbols) or strong (3 consonants) distractor was shown for 0.5 s, followed by the 1.5 s remainder of the retention period. A probe letter was then shown, and the subject was instructed to press the right arrow key if the letter was in the memory set, or the left arrow key if it was not. The probe remained on the screen until the subject responded. Weak and strong distractors were shown in randomised blocks of 20 trials, with 12 blocks for each distractor type (total = 240 trials per condition), allowing participants to anticipate the strength of the distractor within blocks. Distractors were never part of the memory set, and the participants were explicitly told to ignore the distracting stimulus. A short break was allowed between blocks. Prior to the main experiment, participants received a practice block of 20 trials to familiarise themselves with the task.

**Fig. 1.**
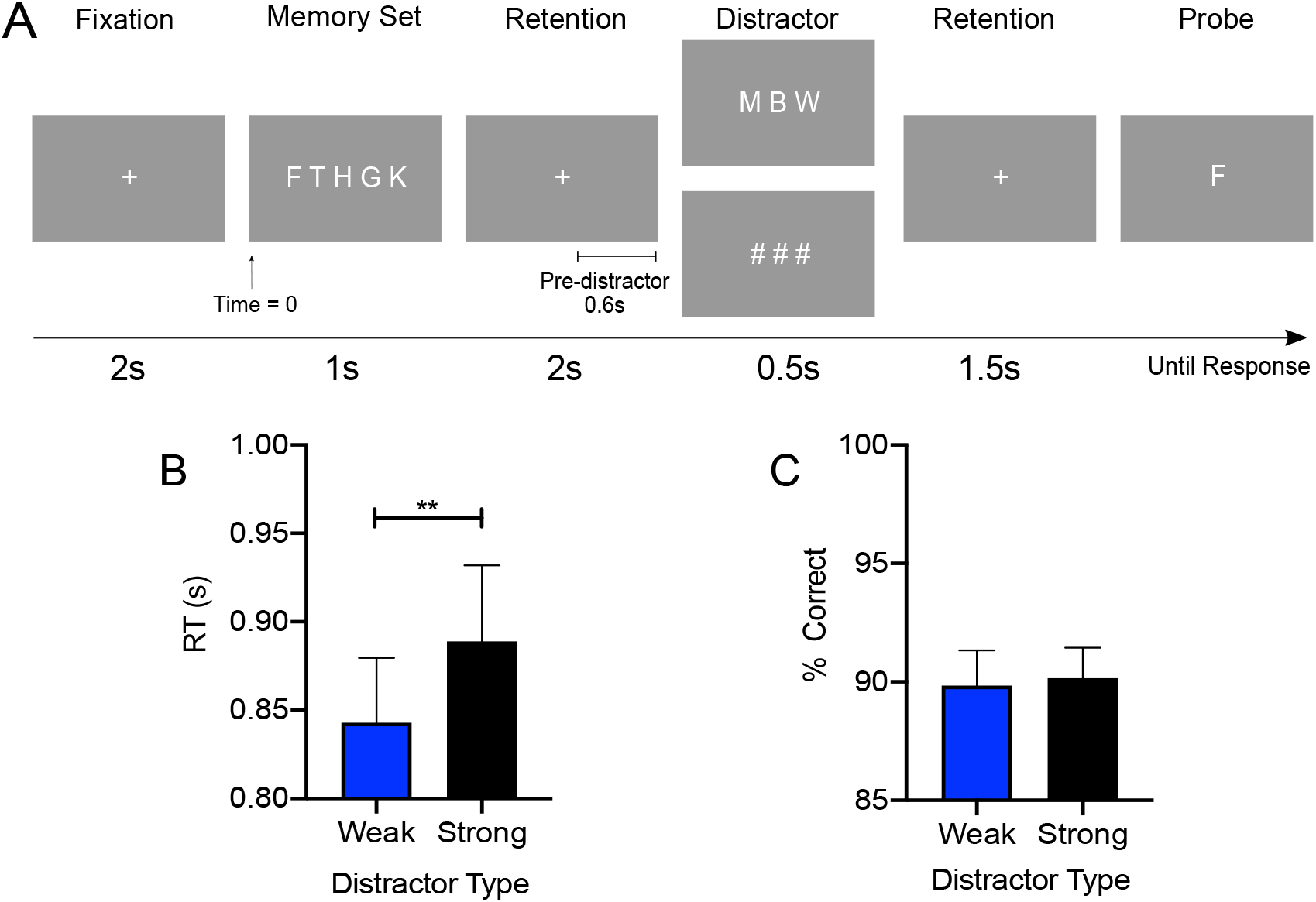
(A) Modified Sternberg task. Each trial contained four stages, including fixation lasting for 2 s, encoding, where a memory set of 5 consonants was displayed for 1 s, a 4 s retention stage where a strong (3 consonants) or weak (3 hash symbols) distractor occurred and persisted for 0.5 s, and a retrieval stage where the subject responded to whether the probe was part of the memory set. (B) Reaction time (RT) for correct responses to the probe and (C) accuracy in response to the probe for strong and weak distractor conditions **p<0.01

### EEG Data Acquisition

EEG data were recorded using a Polybench TMSi EEG system (Twente Medical Systems International B.V, Oldenzaal, The Netherlands) using a 64-channel EEG cap (Waveguard, ANT Neuro, Enschede, The Netherlands). Conductive gel was inserted into each electrode using a blunt-needle syringe in order to reduce impedance to <5kΩ. The ground electrode was located at AFz. Signals were amplified 20x, online filtered (DC-553 Hz), sampled at 2048Hz and referenced to the average of all electrodes. EEG was recorded during each block of 20 trials.

### EEG Pre-Processing

Task EEG data were pre-processed using EEGLAB (Delorme and Makeig, 2004) and custom scripts using MATLAB (R2019, The Mathworks, USA). Due to technical issues, the last block from each participant failed to record. Therefore, to maintain balance of distractor conditions due to the missing block, the final block from the other distractor type was removed prior to pre-processing. Data from each block were then merged into a single file, sorted by distractor type and down sampled to 256 Hz. Noisy and unused channels were removed based on visual inspection, with an average of 1.45 channels removed from each participant (range: 0-4). Data were band-pass (1-100 Hz) and band-stop (48-52 Hz) filtered using a zero-phase fourth order Butterworth filter, then epoched −2 s to 7 s relative to the beginning of the encoding stimulus. Independent component analysis (ICA) was conducted using the FastICA algorithm (Hyvärinen and Oja, 2000), with the ‘symmetric approach’ and ‘tanh’ contrast function, and components corresponding to eye-blinks and persistent scalp muscle activity were removed from the data. Data were then visually inspected to remove any trials contaminated with residual artefacts (e.g. remaining blinks and non-stereotypic artefacts). Missing channels were then interpolated, and data were re-referenced to the common average. Task data were then matched to the epochs, and incorrect trials, as well as trials with outlier RT (defined as >3xSD) were removed, before being split into weak and strong distractor types for spectral analysis.

On average, 180 trials were accepted for final analysis in the weak distractor condition (range 149-203) and 183 in the strong distractor condition (range 155-204).

### Spectral Analysis

FieldTrip toolbox (Oostenveld et al., 2011) was used to analyse task EEG data. Data were converted to the time-frequency domain using a multi-taper transformation based on multiplication in the frequency domain. A time window 3 cycles long was used for each frequency (0.5 Hz steps between 5 to 30 Hz) and time point (20 ms steps), and a Hanning taper was multiplied to the data. Power was calculated for individual trials before averaging for each distractor condition. For analyses across distractor conditions, the data were baseline corrected (dB method) to −0.85 to −0.25 before the onset of the memory set.

### Statistical Analyses

Statistical analyses were performed using R (version 3.4.2) and FieldTrip toolbox (Oostenveld et al., 2011). Paired samples t-tests were used to analyse differences in accuracy and RT between strong and weak distractor conditions. In all tests, a p-value of less than 0.05 was considered statistically significant. Data were presented as mean ± SD in text and mean ± SEM in figures.

Cluster-based permutation tests were used to assess differences in alpha power between baseline and pre-distractor time intervals, and between distractor conditions. Cluster-based permutations control for the type 1 error rate when comparing across multiple channels, frequencies and times (Maris and Oostenveld, 2007). Statistical analyses were restricted to the WM retention period. Clusters were defined as two or more neighbouring electrodes with a p-value <0.05. A permutation distribution was created using the Monte Carlo method (2000 random permutations). A cluster was deemed significant if the p-value of the comparison between the cluster statistic (defined as the cluster with the maximum sum) and the permutation distribution was <0.05 (two-tailed).

## Results

### Behavioural Data

A paired samples t-test revealed that RT was faster for weak distractor trials (0.84 s ± 0.16 s) than strong distractor trials (0.88s ± 0.19 s) (*t*_19_=-3.25, *p*<0.01, *d*=-0.7) (figure 1B). However, we could not find evidence for a difference in accuracy between distractor conditions (*t*_19_=0.38, *p*=0.71) (figure 1C).

### Time Frequency Analysis

Our a-priori hypothesis was that higher alpha power would be present prior to the onset of a strong, relative to a weak distractor. When we examined the time-frequency representation of raw power averaged across both distractor conditions from all participants, the largest alpha power was observed in the 10-13 Hz frequency range, approximately 0.6 s prior to the onset of the distractor. To first determine whether alpha increased in anticipation of the weak and strong distractors, we investigated whether the time period immediately preceding the distractor differed from baseline (−0.85 to −0.25 s before memory set onset). After averaging over the 0.6s preceding the distractor, and over the 10-13 Hz frequency range, cluster-based permutation tests revealed a significant difference between the baseline and pre-distractor time points for weak (*p*=0.015) and strong (*p*=0.023) distractor trials (figure 2A). These differences were most pronounced over the right parietal, parieto-occipital and occipital electrodes. To explore whether differences were present in other frequency bands, we extended out our analyses to include an expanded frequency range (5-30 Hz) and time of interest (1 s before distractor). There were no significant differences between baseline and pre-distractor in weak distractor trials (*p*=0.061) or in strong distractor trials (*p*=0.065).

**Fig. 2.**
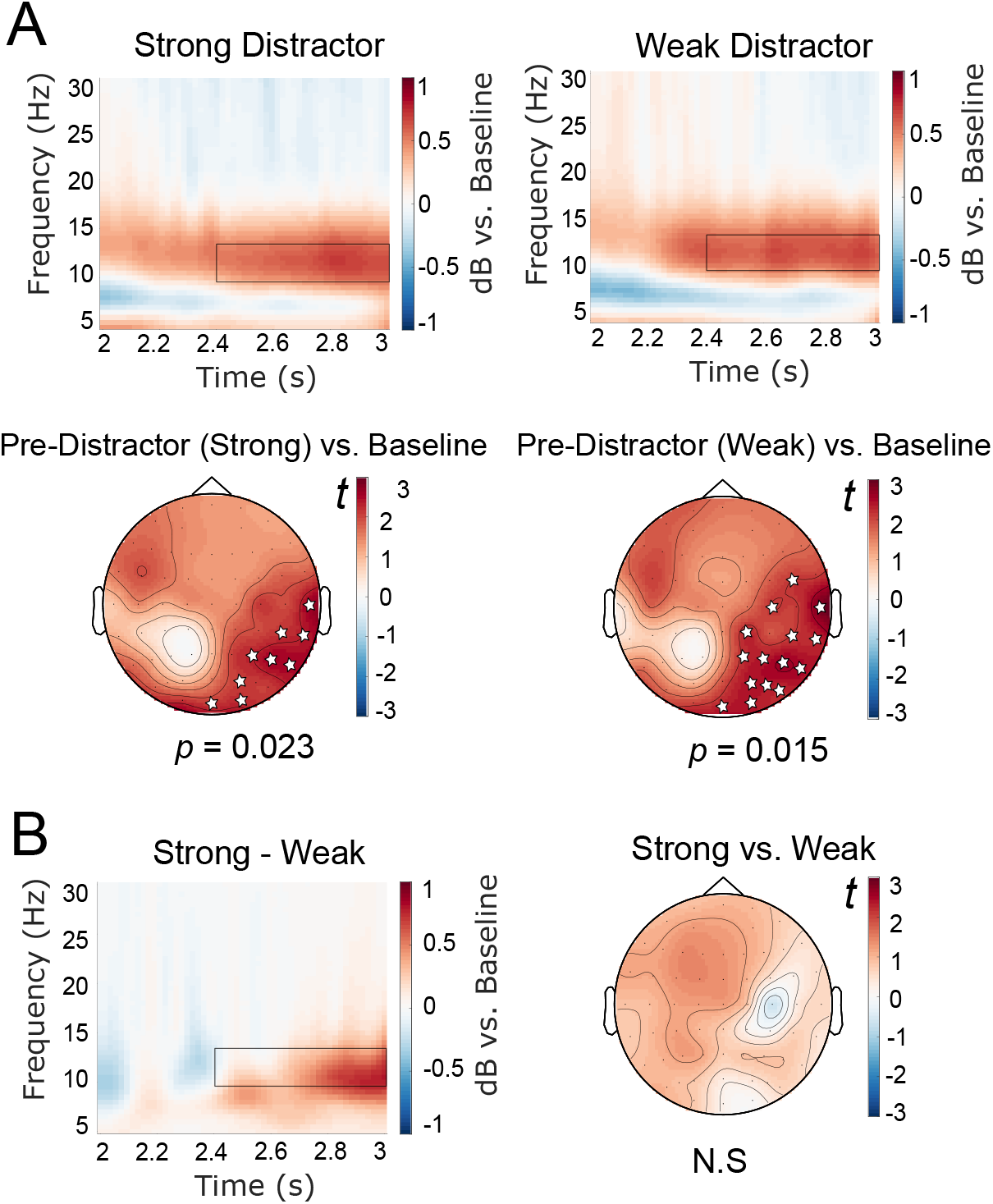
(A) Baseline corrected time frequency representations of power and t-statistics for cluster-based permutation tests for strong (left) and weak (right) distractor conditions. (B) Time frequency representation of power demonstrating the difference between strong and weak distractors prior to distractor onset. The comparison between distractor types yielded no significant differences between conditions. Black boxes in time-frequency plots indicate times and frequencies of interest for cluster-based permutation tests and white stars indicate electrodes in the significant cluster. The onset of the distractor occurred at 3 s

We then examined whether alpha power prior to onset of the distractor differed between strong and weak distractor conditions. Cluster-based permutation tests revealed no significant difference between distractor types in the 10-13 Hz range in the 0.6 s preceding the distractor (no significant clusters). When this analysis was extended out to include the expanded 5-30 Hz frequency range, there were no significant differences found between distractor types at the pre-distractor time point (all *p*>0.09) (figure 2B).

## Discussion

In this study, we investigated the role of alpha power in anticipation of distractors of varying strength during a verbal WM task with a simultaneously presented memory set. While strong distractor trials led to slower RT in response to the probe compared to weak distractor trials, we found no differences in anticipatory alpha power between distractor conditions. However, we found evidence for increases in right parietal, parieto-occipital and occipital alpha power from baseline in anticipation of each distractor type.

Alpha power is often associated with WM retention, although the direction and magnitude of alpha modulation is highly task dependent. In modified Sternberg tasks similar to that in our study, alpha power in visual brain regions has been shown to increase with WM load (Jensen et al., 2002), with this effect interpreted to reflect inhibition of task-irrelevant visual information (Jensen and Mazaheri, 2010). Further evidence for this interpretation came from studies employing distractor paradigms, which found a stronger power increase in occipito-temporal areas in anticipation of strong compared to weak distractors (Bonnefond and Jensen, 2012). Together, these findings suggest an increase in alpha power in posterior brain regions before a distractor reflects inhibition of the visual system, preventing the distractor from interfering with WM retention (Bonnefond and Jensen, 2012; Jensen and Mazaheri, 2010). We find partial support for this hypothesis. While we report an increase in alpha power from baseline compared to the time immediately preceding the distractor, we did not see an increase in alpha power with a stronger distractor, despite slowing of RT. Given that alpha power is shown to increase with WM load, we are unable to conclude whether the increase in alpha power from baseline seen here represents distractor inhibition, or WM retention in general.

Presumably, the ability to anticipate a strong distractor should lead to larger increases in alpha power, as the saliency of the distractor should elicit greater interference on WM retention. Despite the presence of a minor behavioural effect in this study, it is possible that features of the task led to a lack of difference between distractor conditions. In our task, the memory set was presented simultaneously, such that encoding and retention were temporally separated. However, when memory sets are presented sequentially, as in Bonnefond and Jensen (2012), the distractor may be more salient due to its proximity to the encoding period and lack of clear distinction between encoding and retention. Therefore, participants may be more likely to employ a visual strategy to suppress distractors due to the increased task demands, leading to an increase in alpha power to reduce the interference of a strong, relative to a weak distractor. When encoding and retention are temporally separated, increasing alpha power in visual brain regions to inhibit a distractor might not be necessary given a visual strategy may not be employed due to lower task demand. Therefore, we cannot rule out that if our task was more difficult, for example by increasing the number of letters in the memory set or by altering the timing or duration of the distractor, then we might have observed increases in alpha power prior to strong, compared with weak distractor onset.

Our results also partially support the findings of some studies employing lateralised visual WM tasks. For example, in tasks where bilateral arrays of coloured squares are presented, and subjects are cued to memorise either the left or right hemifield and ignore the other (i.e. the un-cued hemifield is a distractor), alpha activity during the retention interval increases in the irrelevant hemifield, with this lateralisation increasing with the amount of distractors in the irrelevant hemifield (Sauseng et al., 2009). Similarly, another study investigating the role of alpha activity in proactive and reactive distractor suppression found that increases in alpha power were present only in proactive filtering (i.e. in anticipation of a distractor, rather than suppressing it once it appears) (Vissers et al., 2016). Conversely, in a visual WM task manipulating the strength of distraction during the retention period, it was found that strong distractors were associated with decreases in alpha power (Schroeder et al., 2018). While we provide some evidence for the role of alpha power in anticipatory distractor filtering similar to that of these studies, it is clear that stimulus presentation methods (simultaneous, sequential or lateralised memory sets), the type of WM stimuli (visual or verbal) and features of the distractor (where, when and how long they are presented for) influence the alpha oscillatory activity involved in WM performance and distractor inhibition.

There are a number of limitations associated with this study. First, the absence of a no-distractor condition makes it unclear whether changes in alpha power during the retention period are due to the distractors or due to WM demands in general. As alpha power has been shown to increase in Sternberg tasks similar to the one used in this study (Jensen et al., 2002; Murphy et al., 2019; Proskovec et al., 2019; Tuladhar et al., 2007; Wang et al., 2016), it is difficult to determine whether the significant difference between baseline and the pre-distractor condition is due to the influence of a load-5 memory set, or due to the distractor. Second, it would have been optimal to compare the two stimulus presentation methods (i.e. both sequential and simultaneous memory sets) in the same experimental session as per Okuhata et al. (2013) to determine the influence of task demands on alpha activity during distractor inhibition. Finally, it is possible that the distractors influenced oscillatory dynamics not investigated in this study. For example, alpha phase has been shown to adjust prior to the distractor (Bonnefond and Jensen, 2012) and phase-amplitude coupling between gamma power and the phase of alpha oscillations results in inhibition of sensory processing (Bonnefond and Jensen, 2015).

Although alpha oscillations have been implicated in distractor inhibition during verbal WM, it is evident that the pattern of alpha activity depends on task features. Here, we show that when the encoding and retention intervals are separated in time, alpha power increases in the lead up to a distractor. However, the strength of a distractor does not influence the magnitude of alpha power increase before distractor onset, despite there being a behavioural effect on WM performance. Future work should now investigate how task demands influence the direction, location and magnitude of alpha oscillations associated with distractor suppression during WM.

## Declarations

### Ethics Approval

This study was approved by The University of Adelaide Human Research Ethics Committee (H-2018-049) and was performed in accordance with the 1964 Helsinki Declaration.

### Consent to Participate

Informed consent was obtained from all participants included in this study.

